# Chromosomal rearrangements and loss of subtelomeric adhesins linked to clade-specific phenotypes in *Candida auris*

**DOI:** 10.1101/754143

**Authors:** José F. Muñoz, Rory M. Welsh, Terrance Shea, Dhwani Batra, Lalitha Gade, Anastasia P. Litvintseva, Christina A. Cuomo

**Affiliations:** Broad Institute of MIT and Harvard, Cambridge, MA, USA; Mycotic Diseases Branch, Centers for Disease Control and Prevention, U.S. Department of Health and Human Services, Atlanta, GA, USA; Division of Scientific Resources, Centers for Disease Control and Prevention, U.S. Department of Health and Human Services, Atlanta, GA, USA

## Abstract

*Candida auris* is an emerging fungal pathogen of rising concern due to its increasing incidence, its ability to cause healthcare-associated outbreaks and antifungal resistance. Genomic analysis revealed that early cases of *C. auris* that were detected contemporaneously were geographically stratified into four major clades. Clade II, also termed East Asian clade, consists of the initial isolates described from cases of ear infection, is less frequently resistant to antifungal drugs and to date, the isolates from this group have not been associated with outbreaks. Here, we generate nearly complete genomes (“telomere-to-telomere”) of an isolate of this clade and of the more widespread Clade IV. By comparing these to genome assemblies of the other two clades, we find that the Clade II genome appears highly rearranged, with 2 inversions and 9 translocations resulting in a substantially different karyotype. In addition, large subtelomeric regions have been lost from 10 of 14 chromosome ends in the Clade II genomes. We find that shorter telomeres and genome instability might be a consequence of a naturally occurring loss-of-function mutation in *DCC1* exclusively found in Clade II isolates, resulting in a hypermutator phenotype. We also determine that deleted subtelomeric regions might be linked to clade-specific adaptation as these regions are enriched in Hyr/Iff-like cell surface proteins, novel candidate cell surface proteins, and an ALS-like adhesin. The presence of these cell surface proteins in the clades responsible for global outbreaks causing invasive infections suggests an explanation for the different phenotypes observed between clades.

**IMPORTANCE:** *Candida auris* was unknown prior to 2009 and since then it has quickly spread around the world, causing outbreaks in healthcare facilities and representing a high fraction of candidemia cases in some regions. The emergence of *C. auris* is a major concern, since it is often multidrug-resistant, easily spread between patients, and causes invasive infections. While isolates from three global clades cause invasive infections, isolates from Clade II primarily cause ear infections and have not been implicated in outbreaks, though cases of Clade II infections have been reported on different continents. Here, we describe genetic differences between Clade II and Clades I, III and IV, including a loss-of-function mutation in a gene associated with telomere length maintenance and genome stability, and the loss of cell wall proteins involved in adhesion and biofilm formation, that may suggest an explanation for the lower virulence and potential for transmission of Clade II isolates.

## OBSERVATION

The emerging fungal pathogen *Candida auris* is increasingly reported as the cause of infections and has become a leading cause of invasive candidiasis in some hospitals, often in severely ill patients (1). *C. auris* isolates are commonly resistant to one or more antifungal drugs and can survive for long periods both in the clinical environment and as a commensal on skin (2). Initially identified in cases of ear infection in Japan and South Korea (3, 4), cases of systemic infection were soon after reported in India, South Africa, and Venezuela (5–7).

Initial genomic analysis of the outbreak identified four major genetic groups corresponding to these geographic regions or Clades I, II, III, and IV (8). Clades I, III, and IV are responsible for the ongoing and difficult to control outbreaks in healthcare facilities worldwide (9). Clade II, also termed the East Asia clade, is predominantly associated with cases of ear infection and appears to be less resistant to antifungals than other clades (10). While a reference genome assembly of a Clade I isolate is commonly used for SNP analyses, the karyotype is known to vary based on whole genome alignment with an assembly of a Clade III isolate (11) and wider analysis of chromosomal sizes (12). To better understand the emergence of this species and phenotypic differences between clades, here we leverage complete reference genomes for isolates from Clades II and IV. We find that the genome of Clade II is highly rearranged and is missing large subtelomeric regions that include candidate cell wall proteins found in all of the other three clades, which may help explain the major difference in clinical presentation between isolates from this clade and those from the global expanding clades causing outbreaks.

### Large chromosomal rearrangements and deletions in *C. auris* Clade II

To investigate genomic differences between clades, we generated complete chromosome scale assemblies for isolates from Clades II and IV. Genome assemblies of B11245 (Clade IV) and B11220 (Clade II) consisted of 7 nuclear contigs corresponding to complete chromosomes with telomeres at both ends, excluding one end that corresponds to rDNA in each assembly and one additional end in B11220 (**Supplementary Table 1**). While the number of chromosomes and total genome size is similar (average 12.3 Mb), chromosome lengths can differ by up to 1.1 Mb in Clade II relative to Clades I, III and IV (**Figure 1a**; **Supplementary Table 1**). By aligning these genomes with those previously published (11), we found evidence of large chromosomal rearrangements (> 10 kb) between the *C. auris* clades (**Figure 1b** to **d**; **Supplementary Table 1**). Notably, the genome of Clade II (B11220) is the most highly rearranged compared to the other three, with two inversions and 9 translocations resulting in large changes in chromosome size (**Figure 1b**). Fewer chromosomal rearrangements were detected in the most distantly related isolate Clade IV (B11245), including two inversions located in chromosome 1 (**Figure 1c**). The Clade III genome (B11221) contains one inversion in chromosome 1 and two translocations in chromosome 1 and 3 ((11); **Figure 1d**). These alterations in karyotype, most dramatically of Clade II, likely serve as a barrier to the production of viable progeny following mating and recombination.

**Figure 1.**
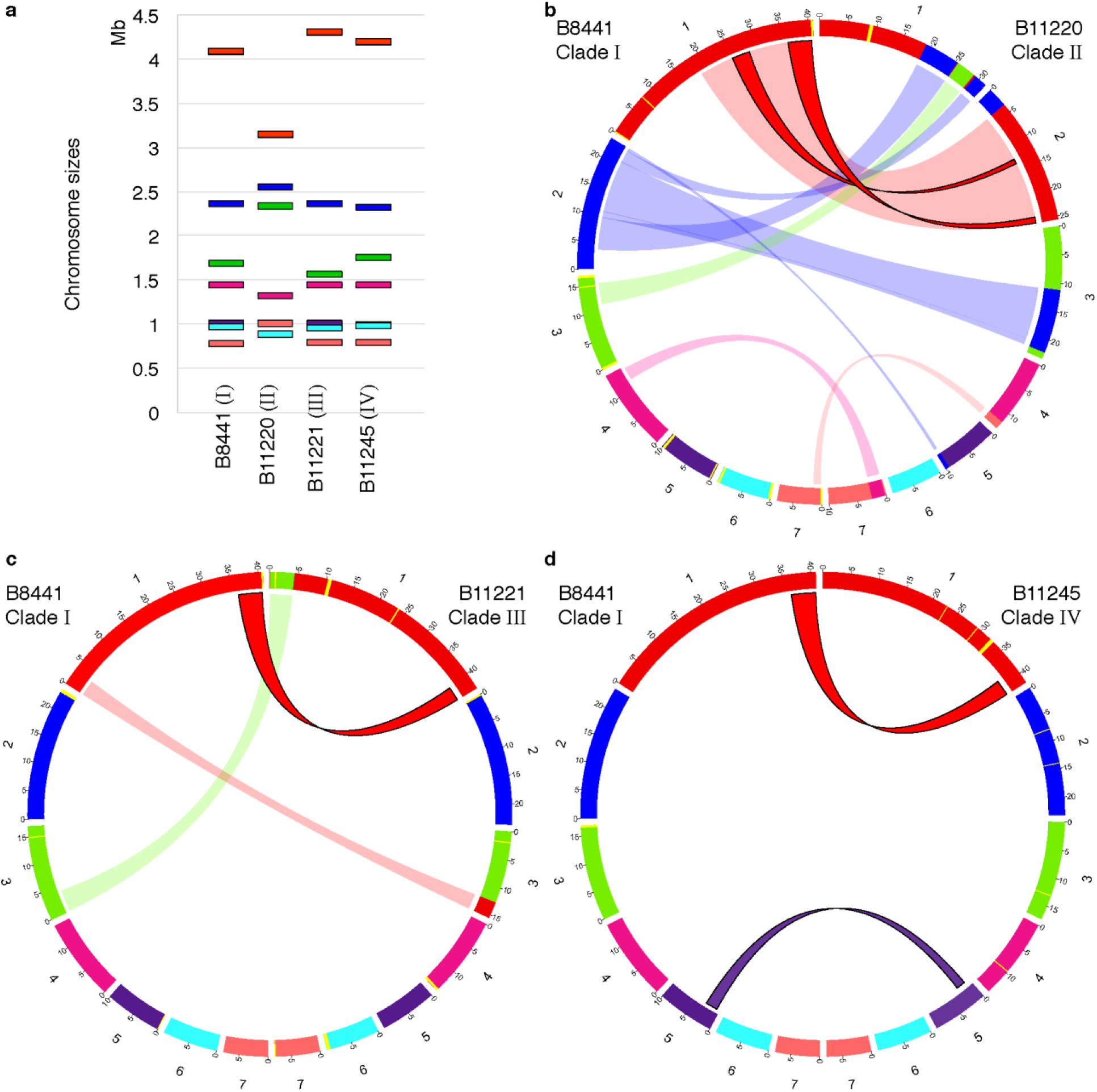
Karyotypic variation and chromosomal rearrangements in *Candida auris*. **(a)** Chromosome sizes are based on complete chromosome scale contigs. **(b-d)** Circos plots showing syntenic chromosomes by color and links for inversions (twisted filled links) and translocations (flat transparent links) using B8441 (Clade I) as reference compared to B11220 (Clade II; **b**), B11221 (Clade III; **c**), and B11245 (Clade IV; **d**). Scaffolds/contigs to chromosome mapping for these genome assemblies is included in **Supplementary Table 4**.

In addition to rearrangements, we identified large genomic regions that were missing in one or more clades, predominantly subtelomeric regions that are absent in Clade II relative to Clades I, III and IV. Comparing the Clade I (B8441) and Clade II (B11220) genomes, we identified 11 large regions (>5 kb) absent in Clade II that encompassed 226 kb and 74 genes; 10 of these 11 regions are subtelomeric in B8441 (**Figure 1b**; **Supplementary Table 2**). Comparing with assemblies of Clade III (B11221) and Clade IV (B11245), we confirmed that these regions were also subtelomeric and only absent in Clade II. These subtelomeric deletions are a common feature of isolates from Clade II, as these regions are also absent in five other Clade II isolates from United States and South Korea, based on aligning whole genome sequence to B8441 (**Supplementary Table 3**). To search for genetic variation that could explain these dramatic changes in genome integrity in Clade II, we examined loss-of-function mutations found exclusively in Clade II isolates (**Supplementary Note**). We found that Clade II isolates have a loss-of-function mutation in *DCC1* (B9J08_000232), specifically a point mutation near the beginning of the protein (amino acid change=Y10*; codon change=taC/taG; **Supplementary Table 3**). As mutations in *DCC1* in *Saccharomyces cerevisiae* result in shorter telomeres (13) and genome instability (14), this suggests that this naturally occurring loss of function mutation in Clade II might contribute to shorter telomeres and genome rearrangements observed in this clade.

### Depleted *Hyr/Iff* and species-specific cell-wall protein families in *C. auris* Clade II

The subtelomeric regions deleted in Clade II likely contribute to the phenotypic differences of this clade, most notably by the loss of fourteen candidate adhesins present in Clades I, III and IV. These include two sets of genes that contain predicted GPI anchors and secretion signals, one set sharing sequence similarity to *C. albicans* adhesins from the *Hyr/Iff* family and a second set of clustered genes only found in *C. auris* and the closely related species *C. haemulonii and C. duobushaemulonii* (**Figure 2a**; **Supplementary Table 2**). The *Hyr/Iff* gene family was previously noted to be the most highly enriched family in pathogenic *Candida* species and has been associated with pathogenicity and virulence (15). Six of eight *Hyr/Iff* proteins found in *C. auris* contain intergenic tandem repeats, which modulate adhesion and virulence (16); five *Hyr/Iff* genes are deleted in Clade II (**Figure 2a**; **Supplementary Table 2**). The second set, *C*. *auris*-specific candidate adhesins, are small proteins with serine/threonine-rich regions (9.0% to 17.3% Serine; 17.5% to 22.8% Threonine), and are tandemly located in subtelomeric regions conserved in Clades I, III and IV, but absent in Clade II (**Figure 2a** and **b**; **Supplementary Table 2**). The subtelomeric location and serine/threonine-rich region are properties shared with *C. glabrata EPA* adhesins (17), and the expansion of *EPA* adhesins is linked to the emergence of the ability to infect humans in the *C. glabrata* lineage (18). Several of the genes in deleted regions of Clade II (three of the *C. auris*-clade specific adhesins, one *HYR-like* gene, and other cell-wall associated proteins (*ALS4*, *CSA1* and *RBR3*)) were induced during developing *C. auris* biofilms ((19); **Supplementary Table 2**), suggesting they play a role in biofilm formation. The loss of adhesins-like genes in Clade II isolates could explain differences in infection and environmental phenotypes between this clade and the others that are more commonly observed.

**Figure 2.**
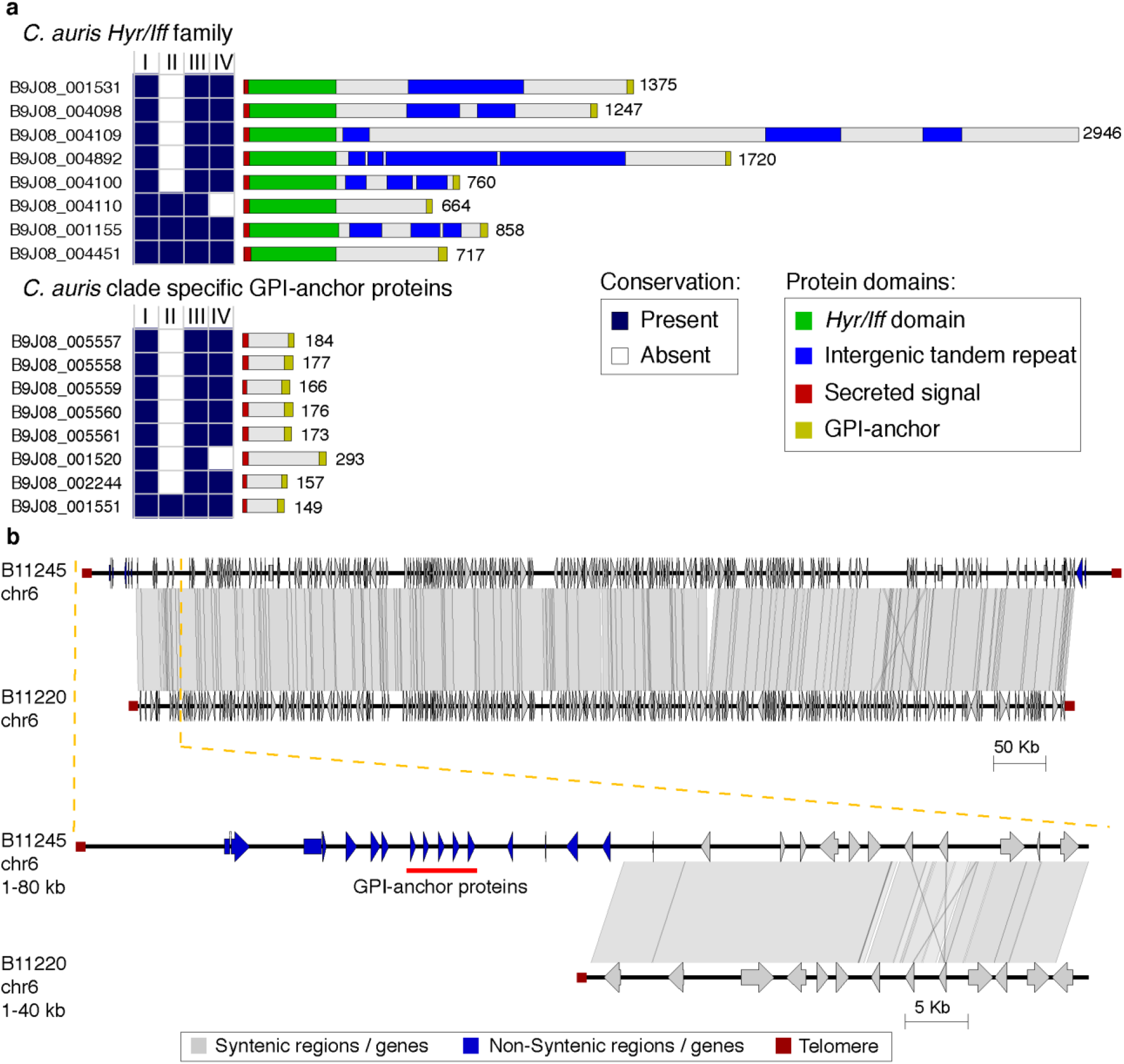
Differences in the repertoire GPI-anchor proteins in *Candida auris*. **(a)** Conserved domains in two clusters of GPI-anchor found in *C. auris* B8441. (top) *Hyr/Iff* GPI-anchor family and (bottom) *C. auris* clade-specific GPI-anchor protein. Conservation across *C. auris* strains representing Clades I, II, III and IV is color-coded indicating whether the gene is present (dark blue) or absent (white). **(b)** Chromosome wide synteny between B11245 (Clade IV) and B11220 (Clade II). Chromosome 6 includes telomeres at both ends in both strains (dark red square). Shared synteny regions based on genome alignment (blastn) are depicted in gray vertical blocks connecting the chromosome regions. Depicted genes in blue and light gray arrows showing the direction of transcription are color-coded according to the location in conserved (gray) or non-conserved (blue) regions. The top comparison corresponds to the entire chromosome 6, and the bottom comparison corresponds to a zoom in of the subtelomeric region depleted in Clade II (B11220), which encompasses *auris*-clade specific adhesins in tandem (red line).

### Phenotypic differences between *C. auris* clades correlate with adherence ability

*Candida auris* can be easily transmitted within health care facilities, accelerated by the ability to persist on plastic surfaces common in health care settings (2), which may be enhanced by its ability to form biofilms. In addition, the ability of *C. auris* to form biofilms is associated with increased resistance to all classes of antifungals (19) and might also enhance *C. auris* capacity to colonize patients’ skin further increasing patient to patient transmission and potentiating outbreaks. These are features of isolates from Clades I, III and IV that are the primary cause of invasive infections and are rarely reported from ear infections (10). The identification of these candidate cell wall proteins that are absent in the highly rearranged genome of Clade II, genomic changes likely due to a natural loss-of-function variant in *DCC1* in Clade II, highlights the major differences that can occur between otherwise closely related isolates of a species. These candidate cell wall proteins are strong candidates to explain the different phenotypic properties of these clades, and further understand the emergence of this species.

## Methods

### DNA Purification

High molecular weight DNA for long read sequencing was obtained using the Epicentre MasterPure yeast DNA purification kit (MPY80200). DNA for Illumina sequencing was extracted using the ZYMO Research ZR Fungal/Bacterial DNA MiniPrep kit.

### Genome assembly

Chromosome-level assemblies for Clade IV (South America) strain B11245 (CDC AR 0386) and Clade II (East Asia) strain B11220 (CDC AR 0381) were generated using an Oxford Nanopore Technology Ligation Sequencing Kit 1D (SQK-LSK108), sequenced on a MinIon Flow Cell R9.4 (FLO-MIN106) and basecalled with Albacore v2.0.2. The total read depth was 88X (21.2 kb N50) for B11245 and 51X (20.6 kb N50) for B11220. Reads were assembled using Canu v1.5 (genomeSize=12000000; stopOnReadQuality=false; correctedErrorRate=0.075) and Flye v 2.4.2 (genome-size=12000000) (20, 21). The most contiguous assembly was obtained with Canu for B11245 and Flye for B11220. A tandem motif (AGACACCACCTA{1,2}GAAA{1,2}CC{1,2}) was identified at contig ends; contig ends missing this motif were aligned to the unassembled contigs and manually extended. Each assembly had five iterations of Illumina read error correction using Pilon v1.12 (22). Assemblies were aligned to each other and to B8441 and B11221 (11) using NUCmer (MUMmer v3.22) (23), and rearrangement sites were manually inspected for read support. Gene annotation was performed using RNA-Seq to improve gene structure predictions as previously described ((11); **Supplementary Note**). The predicted gene number was highly similar across all *C. auris* genomes, totaling 5,328 for B11220 and 5,506 for B11245. GPI anchored proteins were predicted with PredGPI using the general model and selecting proteins with high probability (> 99.90% specificity) (24).

### Genome alignments

Shared synteny regions of at least 10 kb were identified using NUCmer v3.22 (23). Chromosomal rearrangements (translocations, inversions and deletions) were identified from the alignments blocks based on alignment length, chromosome mapping, and orientation. Illumina read alignments were manually inspected in Integrative Genomics Viewer (IGV) v2.3.72 (25) to confirm that the rearrangement junctions are well supported in each assembly. Illumina sequence of 5 additional clade II isolates (B11808, B11809, B12043, B12081, and B14308) were aligned to the B8441 genome using BWA mem v0.7.12 (26) and deleted regions identified using CNVnator v0.3 (1 kb windows; *p*-value < 0.01) (27). Variants were identified between these isolates using GATK v3.7 (28) (**Supplementary Note**).

### Data and resource availability

The whole genome sequence and assemblies of B11220 and B11245 were deposited in NCBI under BioProject PRJNA328792. Illumina sequence of Clade II isolates is available in BioProject PRJNA328792. Isolates are available from the CDC and FDA Antimicrobial Resistance (AR) Isolate Bank, https://www.cdc.gov/drugresistance/resistance-bank/index.html.

### Disclaimer

The use of product names in this manuscript does not imply their endorsement by the US Department of Health and Human Services. The finding and conclusions in this article are those of the authors and do not necessarily represent the views of the Centers for Disease Control and Prevention.

## Acknowledgements

This project has been funded in part with Federal funds from the National Institute of Allergy and Infectious Diseases, National Institutes of Health, Department of Health and Human Services, under award U19AI110818 to the Broad Institute. CAC is a CIFAR fellow in the Fungal Kingdom Program.

